# Role of Stem-Like Cells in Chemotherapy Resistance and Relapse in pediatric T Cell Acute Lymphoblastic Leukemia

**DOI:** 10.1101/2024.06.24.600391

**Authors:** Julia Costea, Kerstin K. Rauwolf, Pietro Zafferani, Tobias Rausch, Anna Mathioudaki, Judith Zaugg, Martin Schrappe, Cornelia Eckert, Gabriele Escherich, Jean P. Bourquin, Beat Bornhauser, Andreas E. Kulozik, Jan O. Korbel

**Affiliations:** Molecular Medicine Partnership Unit (MMPU), EMBL and Medical Faculty of the University of Heidelberg, Heidelberg, Germany; European Molecular Biology Laboratory (EMBL), Genome Biology Unit, Heidelberg, Germany; Faculty of Biosciences, Heidelberg University, Heidelberg, Germany; Division of Pediatric Oncology, University Children’s Hospital, Zürich, Switzerland; Molecular Biosciences/Cancer Biology Program, Heidelberg University and German Cancer Research Center (DKFZ), Heidelberg, Germany; European Molecular Biology Laboratory (EMBL), Genomics Core Facility, Heidelberg, Germany; European Molecular Biology Laboratory, Structural and Computational Biology Unit, Heidelberg, Germany; Department of Pediatrics, University Hospital Schleswig-Holstein, Campus Kiel, Kiel, Germany; Department of Pediatric Oncology/Hematology, Charité Universitätsmedizin Berlin, Berlin, Germany; German Cancer Consortium (DKTK), and German Cancer Research Center (DKFZ), Heidelberg, Germany; Clinic of Pediatric Hematology and Oncology, University Medical Center Hamburg-Eppendorf, Hamburg, Germany; Department of Pediatric Oncology, Hematology, and Immunology, University of Heidelberg, Heidelberg, Germany; Hopp Children’s Cancer Center (KiTZ) Heidelberg, Heidelberg, Germany; Clinical Cooperation Unit Pediatric Leukemia, German Cancer Research Center (DKFZ), Heidelberg, Germany; Bridging Research Division on Mechanisms of Genomic Variation and Data Science, German Cancer Research Center (DKFZ), Heidelberg, Germany

**Author notes:** Correspondence: Andreas E. Kulozik, Department of Pediatric Oncology, Hematology and Immunology, University Hospital Heidelberg, Im Neuenheimer Feld 430, 69120 Heidelberg, Germany,; and Jan O. Korbel, Genome Biology Unit, European Molecular Biology Laboratory, Meyerhofstrasse 1, 69117 Heidelberg, Germany.

## Abstract

T cell acute lymphoblastic leukemia (T-ALL) is an aggressive leukemia predominantly affecting adolescents and young adults. T-ALL relapses are characterized by chemotherapy resistance, cellular heterogeneity and dismal outcome. To gain a deeper understanding of the cellular heterogeneity and mechanisms driving relapse, we conducted single-cell full-length RNA sequencing of 13 matched pediatric T-ALL patient-derived xenografts (PDX) samples obtained at initial diagnosis and relapse, generating the to date most comprehensive longitudinal single cell study in paired T-ALL samples, along with 5 non-relapsing PDX samples collected at initial diagnosis. This dataset identifies considerable transcriptomic diversity among individual T-ALL cell populations. Notably however, 11 of the 18 patients exhibit a small T-ALL cell subpopulation with a shared set of gene regulatory networks characterized by a common set of active regulons, expression patterns and splice isoforms. This profile involves the upregulation of a stem-like cell signature with enrichment of cell adhesion, and inhibition of the cell cycle and metabolic activity. Comprehensive investigations of these networks identify transcripts enforcing drug resistance through NF-κB expression and TGF-β signaling and of anti-apoptotic T cell signaling. Longitudinal monitoring of these stem-like cells demonstrates this subpopulation to account for only a small proportion of leukemia cells initially with a substantial expansion at relapse suggesting resistance to first line therapy. Chemotherapy resistance is functionally corroborated through *in vitro* and *in vivo* drug testing. We thus report the discovery of inherently treatment-resistant stem-like T-ALL cell populations underscoring the potential for devising future therapeutic strategies aimed at targeting stemness pathways in pediatric T-ALL.

**Key Points:**

- Single-cell full-length RNA sequencing reveals dormant, treatment-resistant cells expanding upon T cell acute lymphoblastic leukemia relapse.
- Stem-like T-ALL cells exhibit a shared gene regulatory network conferring chemotherapy resistance and relapse.

## Introduction

ALL is the prevalent form of pediatric leukemia affecting approximately 1 out of 3 of pediatric cancer patients. T-ALL comprises ∼15 % of pediatric ALL and can be cured in ∼80% of affected patients^1^. However, the outcome in relapsing patients is dismal with a prognosis of merely 20% survival^2^. Although the acquisition of mutations and changes in the expression of genes with roles in pharmaco-resistance, as well as in epigenetic regulation have previously been described^3–7^, a unifying mechanism explaining the development of relapse remains unknown.

A differential analysis of disease progression and relapse in T-ALL has been facilitated by the identification of two relapse categories: type-1, originating from the major clone of the initial disease and type-2, originating from a minor ancestral clone^3,5^. Type-1 relapses are characterized by frequent activation of the IL7 receptor pathway, whereas type-2 relapses are driven by activation of the transcription factor TAL1 and are enriched in mutations of cancer predisposing genes^5^. Considering that a substantial proportion of relapses originate from a minor diagnostic subclone, bulk genetic and transcriptomic analyses are insufficient for characterizing treatment resistance acquisition. To comprehensively investigate the heterogeneity of T-ALL in single cells and gain a better understanding of mechanisms driving T-ALL relapse, we therefore conducted single-cell full-length total RNA sequencing with VASA-seq^8^, following the evolution of treatment-resistant clones from initial diagnosis to relapse in the same patient. Our analysis in 18 T-ALL patients identifies a cell population that converges at a gene-regulatory network revealing a common dormant stem-like cell phenotype with resistance to chemotherapy in functional assays.

## Methods

### Patients

The primary cells were obtained from patients recruited in ALL-BFM 2000, ALL-BFM-2009, CoALL97, CoALL03, CoALL09, and ALL-REZ BFM 2002 trials. Patients’ clinical characteristics have been described previously^5^.

Clinical trials from which samples were used in this analysis had previously received approval from the relevant institutional review boards or ethics committees. Written informed consent had been obtained from all the patients or legal guardians, and the experiments conformed to the principles set out in the WMA Declaration of Helsinki and the Department of Health and Human Services Belmont Report.

### Patient-derived xenografts (PDXs)

We maintained T-ALL patient cells as PDXs as previously described^9^. *In vivo* experiments were approved by the veterinary office of the Canton of Zurich, in compliance with ethical regulations for animal research.

### *In vitro* treatment with Cytarabine

MSCs were seeded in 24-well plates at a concentration of 500.000 cells per well in 1 ml AIM V medium. After 24 hours, T-ALL cells were added at a concentration of 1.5 million cells per well in 1 ml AIM V. Cytarabine (MedChemExpress, HY-13605) or DMSO (vector) as control was added after an additional 24 hours at a concentration of 1 μM. After 72 hours cells were trypsinized, collected and frozen in 90% FBS/10% DMSO.

### *In vivo* drug treatment

Mice with a human engraftment above 5% measured through flow cytometry using mCD45 (eFluor 450, eBiosciences, 1:100), hCD7 (PE, eBiosciences, 1:25) and hCD45 (Alexa Fluor 647, BioLegend, 1:25) were started on treatment which consisted of 0.50 mg kg−1 of vincristine (1 mg ml−1 injection, Teva Pharma) administered weekly intraperitoneally, 10.5 mg kg−1 of dexamethasone (4mg ml-1, Mepha Pharma AG) administered daily intraperitoneally, and 2 mg kg−1 of doxorubicin (2mg ml-1, Sandoz) administered weekly intravenously. After 4 weeks of treatment the leukemia cells were collected by flushing the bone marrow.

### Single cell RNA sequencing

For VASA-seq, mCD45-DAPI- were sorted as single cells into cooled 384-well plates and libraries were prepared by the company according to the VASA-seq protocol^8^ and sequenced on the Nova Seq X Plus (10B - 100 cycle, paired-end, 150.000 reads/cell).

For 10x Genomics bulk of mCD45-DAPI- cells were sorted and samples were processed immediately according to standard 10x Genomics Chromium 3′ (v.3.1 Chemistry) protocol. Libraries were sequenced on the NovaSeq 6000 S2 v1.5 (100 cycles) flowcell S2 (Surface: 3.3-4.1 BIO reads).

### Alternative Splicing (AS) Analysis

The pre-processing of the raw FASTQ files from VASA-Seq was performed via a custom Snakemake^10^ pipeline.

### Data Sharing Statement

For original data, please contact.

See supplemental Methods for further details.

## Results

### Full-length single-cell RNA sequencing of a diagnostic/relapse T-ALL sample pair reveals the expansion of a subclone with a stem-like cell phenotype

To investigate differences in the subclonal architecture of T-ALL during disease progression, we first conducted VASA-seq on initial and relapse samples of our index patient (P2) (Figure 1), generating data for 1,231 single cell full-length transcriptomes altogether (initial: 653; relapse: 578). Uniform Manifold Approximation and Projection (UMAP) analysis identified a predominant cell population (cluster 0, 98.62% of the total population) at initial diagnosis which decreases to 3.27% in relapse. Conversely, an initial minor cell population (cluster 2) expands from 1.37% at diagnosis to 26.47% at relapse suggesting treatment resistance and clonal selection. In addition, the patient acquired a second relapse-specific cell population that has not been detect at initial disease (cluster 1, 70.24 %) (Figure 1b-e).

**Figure 1:**
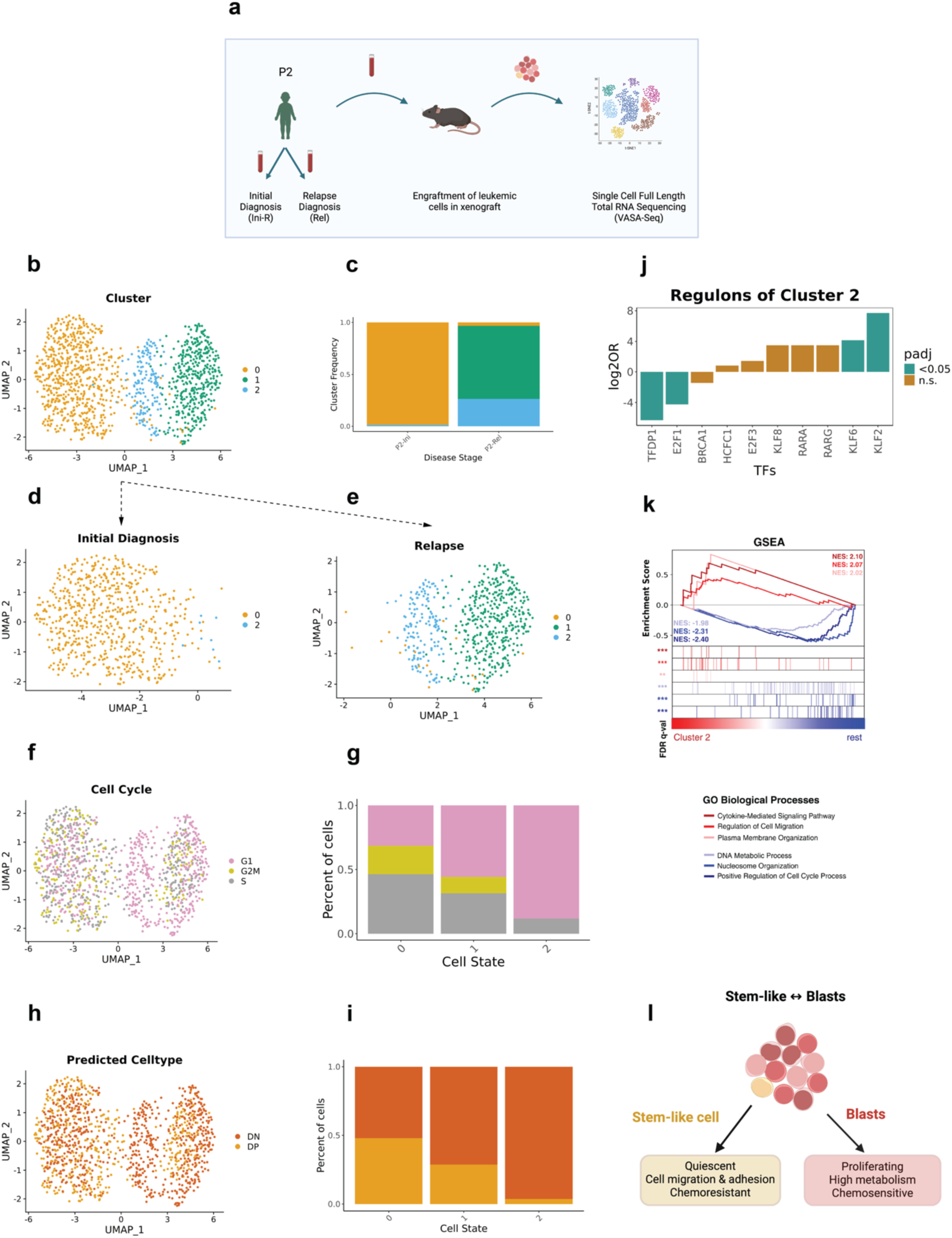
Expansion of a stem cell-like immature dormant subclone during the development of relapse in a child with T-ALL. a) Scheme of the experimental procedure. Created with BioRender.com. The samples have been obtained from patient P2 at the time of initial diagnosis (Ini) and relapse (Rel) and were engrafted in immunodeficient mice. T-ALL cells have been extracted from the PDX mice and analyzed by VASA-seq. b), d), e): UMAP illustration of the clonal composition at initial diagnosis (d) and relapse (e) and combined (b). c) stacked barplot illustrates the cellular composition at initial diagnosis and relapse. f), g): distribution of cells in different cell cycling phases as inferred by the transcriptional profile visualized as UMAP (f) and stacked barplot (g). h), i): distribution of predicted cell types after mapping cells onto a human thymic reference dataset^11^ visualized by UMAP (h) and stacked barplot (i). j) Average enrichment and depletion of regulons in cluster 2 identified by SCENIC (log2FC > 0.25, padj <0.05). k) Gene set enrichment analysis (GSEA) plot of GO biological processes showing significant enrichments of cluster 2 (red) vs rest (blue). l) Illustration of stem-like cell features vs features of the two major clones. Created with BioRender.com.

Subsequent assessment of these clusters in terms of cell cycle and developmental stage (Suppl. Methods) revealed a significant higher fraction of cells in the G1 phase of the cell cycle for cluster 2 (88.27 % in G1) compared to the other clusters (permutation test: FDR <0.05, log2FD >0.25) (Figure 1f-g; Supplemental Figure 1e-g). When mapped onto a thymic single cell reference^11^, cluster 2 significantly differs from cluster 0 and cluster 1 (permutation test: FDR <0.05, log2FD >0.25, Supplemental Figure 1a-c) by almost completely resembling a more immature double negative (DN) phenotype (96.30% DN, 3.70% DP). In contrast, the transcriptional profiles of the other clusters resemble a mixture of the DN and the double positive (DP) phenotype (cluster 0: 47.96% DN, 52.04% DP, cluster 1: 71.18% DN, 28.81% DP) (Figure 2h-i).

**Figure 2:**
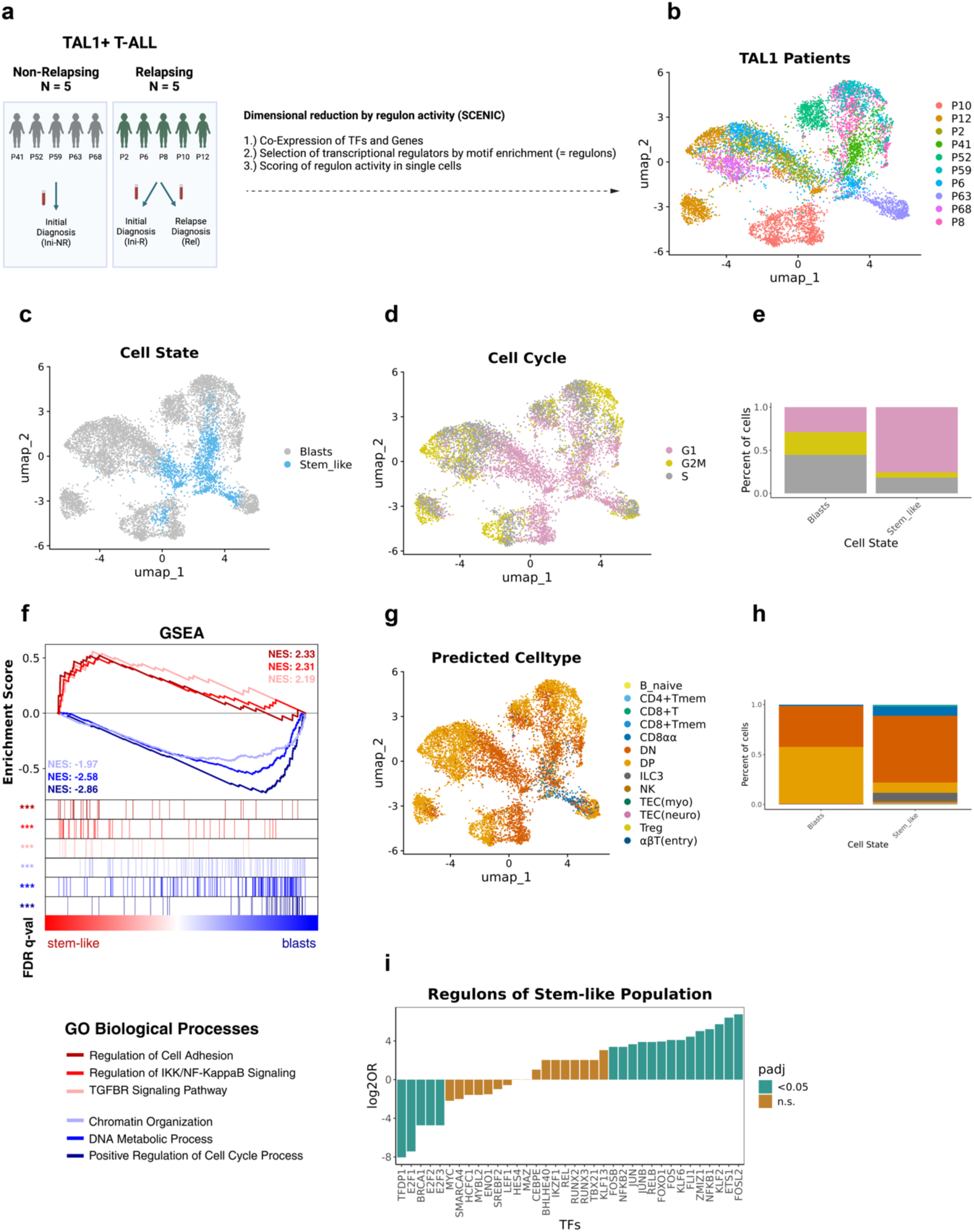
SCENIC analysis reveals a shared regulon activity in small subclones in TAL1- driven T-ALLs with stem cell-like properties. a) Illustration of a TAL1-driven T-ALL patient cohort consisting of 5 non-relapsing patients (Ini-NR, grey), and 5 type-2 relapsing patients (dark green). Samples have been taken during initial diagnosis (Ini-R) and relapse (Rel). Created with BioRender.com. b), c), d), g): UMAP reduction by SCENIC regulon activity. b) distribution of TAL1-driven T-ALLs. c) assignment of stem-like cells and highly proliferative population in TAL1 cohort. d) distribution of cells in different cell cycling phases inferred by transcriptional profile. g) distribution of predicted cell types after mapping cells onto a human thymic reference dataset^11^. e), h) Stacked barplots comparing distribution in stem-like cells and other leukemic blasts. e) predicted distribution of cells in different cell cycling phases. h) distribution of predicted cell types. f) Gene set enrichment analysis (GSEA) plot of GO biological processes showing significant enrichments of stem-like cells (red) vs other leukemic blasts (blue). i) Average enrichment and depletion of regulons of stem-like cells in all patients (log2FC > 0.25, padj <0.05 highlighted in green, non-significant displayed in brown).

In a next step, we investigated functional differences of these dormant, immature cells of cluster 2 compared to leukemic cells of the other clusters utilizing gene set enrichment analysis (GSEA; Suppl. Methods) (Figure 1k). A significant enrichment of cell migration (FDR <0.05, NES 2.07) and plasma membrane organization (FDR <0.1, NES 2.02) alongside a reduction of cell cycling (FDR <0.05, NES −2.40) and metabolic processes (FDR <0.05, NES −1.98) as well as nucleosome organization (FDR <0.05, NES −2.31) resemble characteristics which have been previously noted in stem-like cells^12,13^ (Figure 1l).

Taken together, VASA-seq analysis in this index patient revealed a small cell population at initial diagnosis of cells exhibiting dormant stem cell-like properties, which is substantially expanded during relapse.

### SCENIC network analysis identifies a uniform and shared population among TAL1 driven T-ALLs

We next set out to verify these findings in a group of 9 additional TAL1 driven T-ALLs which either did not (5 patients) or did (4 patients) develop a relapse following first-line treatment thus increasing the total number of patients to 10 (Figure 2a-b). We performed VASA-seq, generating data for 7,929 additional (total: 9,160) single-cell full length transcriptomes. To analyze each single cell with respect to transcriptomic state and regulatory networks, we performed data analysis using the SCENIC single-cell regulatory network inference and clustering workflow (Suppl. Methods)^14,15^. UMAP analysis using regulon activities inferred by SCENIC clustered the majority of cells by individual patients (Figure 2c). However, a small proportion of cells from different patients was grouping closely, indicative for a cell subpopulation driven by a set of common regulons.

As previously seen in cluster 2 of our index patient, a predominant fraction of cells in this subpopulation were arrested in the G1 phase (75.75%), with the remaining 24.25% progressing through S/G2M phases, in stark contrast to the heterogenous predominant subpopulations of the individual patients, which display significantly higher rates of cell division (28.90% G1, 71.09% in S/G2M, permutation test: FDR <0.05, log2FD >0.25) (Figure 2d-e; Supplemental Figure 1h). These data imply the existence of a small and distinct cell population shared across different T-ALL patients, which is characterized by low cell division and which we characterized in further detail in the following.

Notably, when assessing these single cells in terms of their developmental stage, it became apparent that the transcription profile of this quiescent cell population predominantly resembles immature double negative (DN) cells (66.90%, 10.25% DP). By contrast, the highly proliferative population mostly comprises cells with a transcriptional profile resembling double positives (DP) (40.99% DN, 56.98% DP, permutation test: FDR <0.05, log2FD >0.25) (Figure 2g-h; Supplemental Figure 1d).

We next delved deeper into the single cell transcriptomic data to investigate specific pathways active in this dormant cell population. Performing GSEA revealed a strong enrichment in cell adhesion (NES 2.33, FDR <0.05), as well as NF-κB (NES 2.31, FDR < 0.05) and TGF-β signaling (NES 2.19, FDR <0.05) which are known to enforce drug resistance^16,17^ (Figure 2f). Additionally, unlike the cell population with a transcriptional profile of rapidly proliferating cells, the cells with a dormancy phenotype showed a marked downregulation of metabolic processes (−2.86, FDR <0.05, FDR), a common characteristic of stem-like cells^13^. In addition, we find significant enrichment of pathways linked to T-cell quiescence (NES 2.07, FDR <0.01)^18^ as well as clinical correlates of treatment resistance such as prednisone resistance (NES 1.94, FDR <0.01)^19^ and persistent minimal residual disease (NES 1.78, FDR <0.05) (MRD)^20^ (Supplemental Figure 2e-g) in the dormant population. We reasoned that presence of a common cell subpopulation characterized by stem-cell like properties could be of potential relevance for T-ALL therapy resistance, since chemotherapy preferentially targets dividing cells.

### Identification of RNA markers and active regulons of the stem-like cell population

To gain further insights into the transcriptional program of these cells, we next performed comprehensive differential expression analyses. We find that the stem-like cells, unlike the other cell subpopulations, are characterized by a similar RNA expression pattern including several genes that have been recently reported to play a role in stem-like cells in a murine T- ALL model, e.g., CD44^21^, RORA, CD226, CD52^12^ (Supplemental Figure 2a, Supplemental Table 2). Notably, we observed markers that have been suggested to play a role in treatment resistance but were not described in the concept of stemness in T-ALL, e.g., ADGRE5 (CD97)^22^, AHNAK^23^, CD27^24^, COL6A2^25^, ITGB7^22^, several members of the S100 family^26^ and the anti-apoptotic proteins BCL-6, BCL-2 and MCL1. Remarkably, NOTCH1 is downregulated in the stem-like cell population.

We next utilized SCENIC allowing us to identify those TFs that drive the expression of most of the genes in the cluster and therefore are likely to play a particularly relevant biological role (Figure 2I, Supplemental Table 3). We identified five TFs that were significantly less active in the stem-like cells (Fisher’s exact test, padj <0.05), all of which are involved in cell cycling (TFDP1, E2F1, BRCA1, E2F2, E2F3), in further support of the quiescent phenotype of these cells. The 14 TFs with significantly enriched activity in the stem-like cell population (Fisher’s exact test, padj < 0.05) are grouped into six different TF families – KLF, AP1, NF- κB, ETS, FOXO and PIAS family – all of which are key regulators of the hematopoietic development. Notably, KLF2, one of the significantly enriched TFs, is the main driver of dormancy, migration and anti-apoptotic signaling in T cells^27–29^.

### SCENIC analysis enables the development of a stemness score that identifies differential proportions of stem-like cells in T-ALLs driven by different transcription factors

We next investigated whether these stem-like cells are unique to the TAL1 subgroup or are also present in the NKX2, HOXA and TLX1/3 subgroups of T-ALL. Considering that TAL1 driven T-ALLs most frequently relapse as type-2 and the other subgroups frequently relapse as type-1^3,5^, we also considered the relapse-type in our analysis. We thus expanded our patient cohort, now encompassing VASA-seq samples from 13 relapsing (5 type-1, 8 type-2) and five non-relapsing patients of whom 11 were TAL1-, three TLX1/3-, two NKX2-, and two HOXA-driven. While 10 out of the 11 TAL1 driven patients showed a type-2 relapse only one experienced a type-1 relapse (P11). Of the 7 patients with the less common drivers four experienced a type-1 and three a type-2 relapse (Figure 3a).

**Figure 3:**
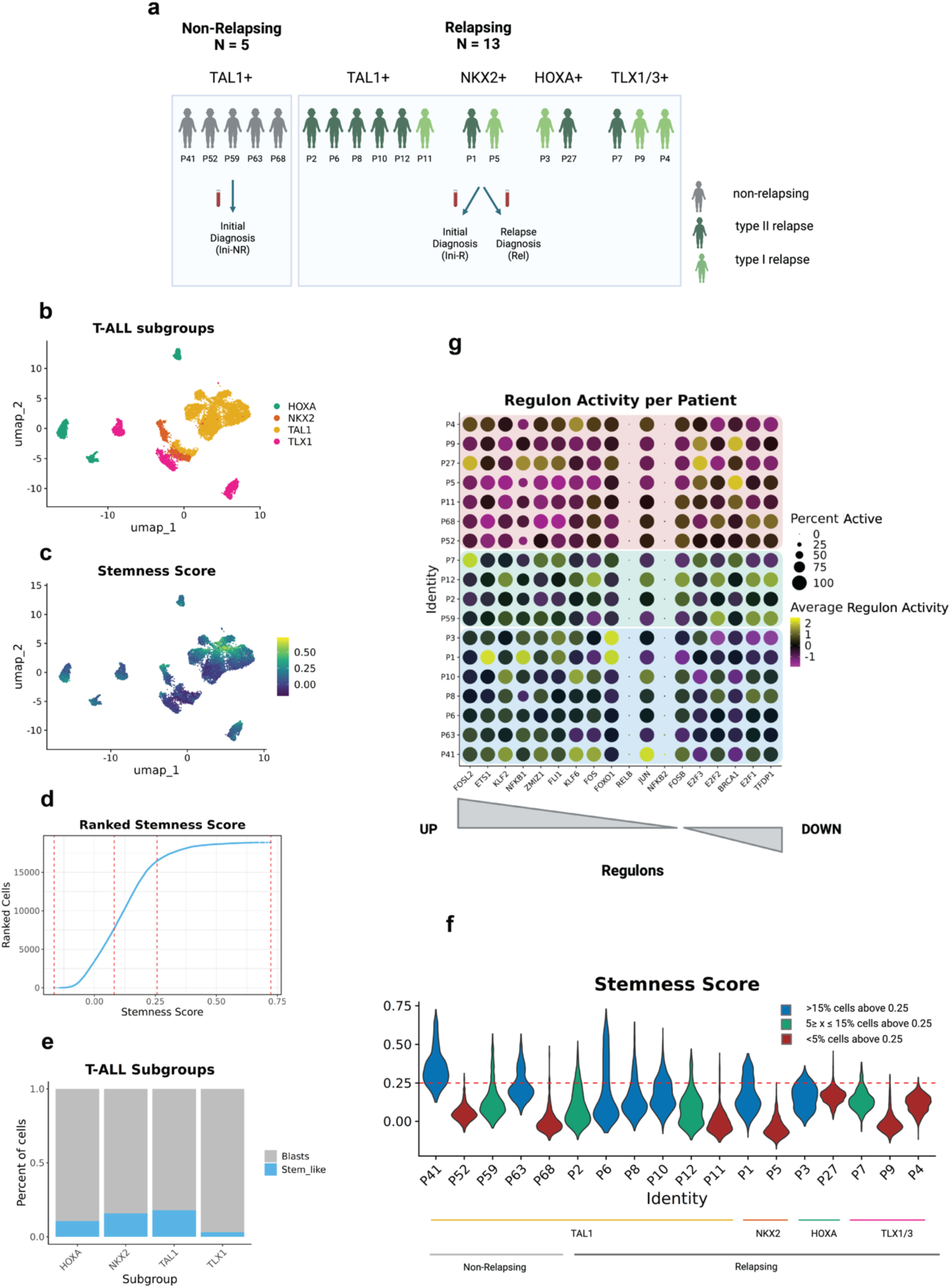
Stem-like cells are more prevalent in TAL1-driven T-ALL than in the other subgroups. a) Illustration of patient cohort consisting of 5 non-relapsing patients (grey), 8 type-2 relapsing patients (dark green) and 5 type-1 relapsing patients (light green). Samples reflect four different T-ALL subgroups: TAL1 (11x), NKX2 (2x), HOXA(2x), TLX1/3 (3x). Samples have been obtained either at initial diagnosis (Ini-NR and Ini-R) or at relapse (Rel). Created with BioRender.com b), c): UMAP reduction of 18 patients using SCENIC regulon activity. b) distribution of T-ALL subgroups. c) Stemness Score is calculated using previously defined expression markers of TAL1 stem-like cell population (log2FC > 0.5, padj <0.05) (Suppl. Methods). d) Cells of all patients ranked by stemness score. Score is calculated as described in Figure 3c-d). Natural breaks in the scores were identified by getJenksBreaks (k=4). Cells with stemness score > 0.25 are considered stem-like cells in the follow-up analyses. e) Stacked barplot displays fractions of stem-like cells vs other leukemic blasts per T-ALL subgroup defined by a stemness score > 0.25. f) Stemness score of cells in individual patients. Score is calculated as described in c,d). Red dashed line reflects the threshold used for the definition of stem-like cells (0.25) that is based on the Jenks natural break classification. Patients are grouped in three different categories based on the proportion of stem-like cells: Blue > 15 %, green 5-15 %, red < 5 %. g) Activity of previously defined regulons of TAL1 stem-like cells calculated in individual patients. Blue, green and red background color corresponds to the different stem-like categories defined in f). Partially created with BioRender.com

Data generation by VASA-seq applied to these patient samples resulted in 9,718 additional (total: 18,878) sequenced full-length transcriptomes. Utilizing the SCENIC output to perform dimensional reduction on all samples together revealed a higher degree of heterogeneity among individual patient samples of the other subgroups compared to the TAL1 subgroup (Figure 3b; Supplemental Figure 2d). To identify stem-like cells in the dimensional space, we used the set of marker genes (log2FC > 0.5, padj <0.05) of the previously defined stem-like cell population of the TAL1 samples to create a signature of ‘T-ALL stemness’ (Suppl. Methods), and scored each cell accordingly (Figure 3c-d; Suppl. Methods). Within the TAL1 subgroup, these data further corroborate the finding that the stem-like cells are more homogenous than the major leukemic cell population. The sample of the type-1 relapsing patient P11, which neither clusters with the other TAL1 samples nor shows an enrichment in stemness, provides a notable exception to this. Remarkably, we do see an enrichment in the stemness score in the other T-ALL subgroups suggesting that stem-like cells are a common phenomenon in T-ALL and not exclusive for TAL1 patients. Nonetheless, the proportion of stem-like cells significantly differs among the subgroups, with TAL1 showing the highest and TLX1/3 patients the lowest (permutation test: FDR <0.05, log2FD >0.25) Supplemental Figure 3a-f). We deconvoluted the stemness score by cells of individual patients (Figure 3f) and categorized each T-ALL based on its proportion of stem-like cells into one of three categories. The group with the highest proportion (>15% of the total population, shown in blue) consists of 5/11 TAL-1 patients, 1/2 NKX2 patients, 1/2 HOXA patients and 0/3 TLX1 patients. Analyzing the activity of the previously identified regulons (Figure 2i) in individual patients confirmed that these T-ALLs are best in reflecting the activity of up- and downregulated TFs while the patients with the lowest proportion of stem-like cells (<5%, shown in red) (3/11 TAL1, 1/2 NKX2, 1/2 HOXA and 2/3 TLX1) show the opposite trend (Figure 3g). Of note, 7/8 relapsing patients with a stem-like cell population >5% had a type-2 relapse and only one patient had a type-1 relapse. Among the TAL1 patients, 3/5 non-relapsing patients and 5/6 relapsing patients show a stem-like cell population with a frequency of >5% (shown in green and blue) suggesting that the presence of a stem-like cell population does not *per se* predict the risk of relapse.

### Common patterns of alternative splicing (AS) enriched in stem-like cells

Over the past decade, increasing evidence emerged that AS is highly correlated with ALL treatment resistance ^30–32^, and acts as a potential driver in pediatric cancers^33^. Harnessing the full-length transcriptome dataset provided through VASA-seq, we investigated splicing usage differences in the T-ALL patient cohort under study with a particular focus on AS in the stem-like cell population, using a previously described computational workflow^8^ to identify AS patterns in single cells (Suppl. Methods, Supplemental Figure 4a-b). Analysis of the different splicing event types revealed the prevalence of four AS event types enriched in the stem-like cell population: intron retention, core exon skipping, usage of an alternative donor, and use of an alternative acceptor site (Figure 4b). The number of total AS events largely varies among individual patients (28- 356) (Supplemental Table 5).

**Figure 4:**
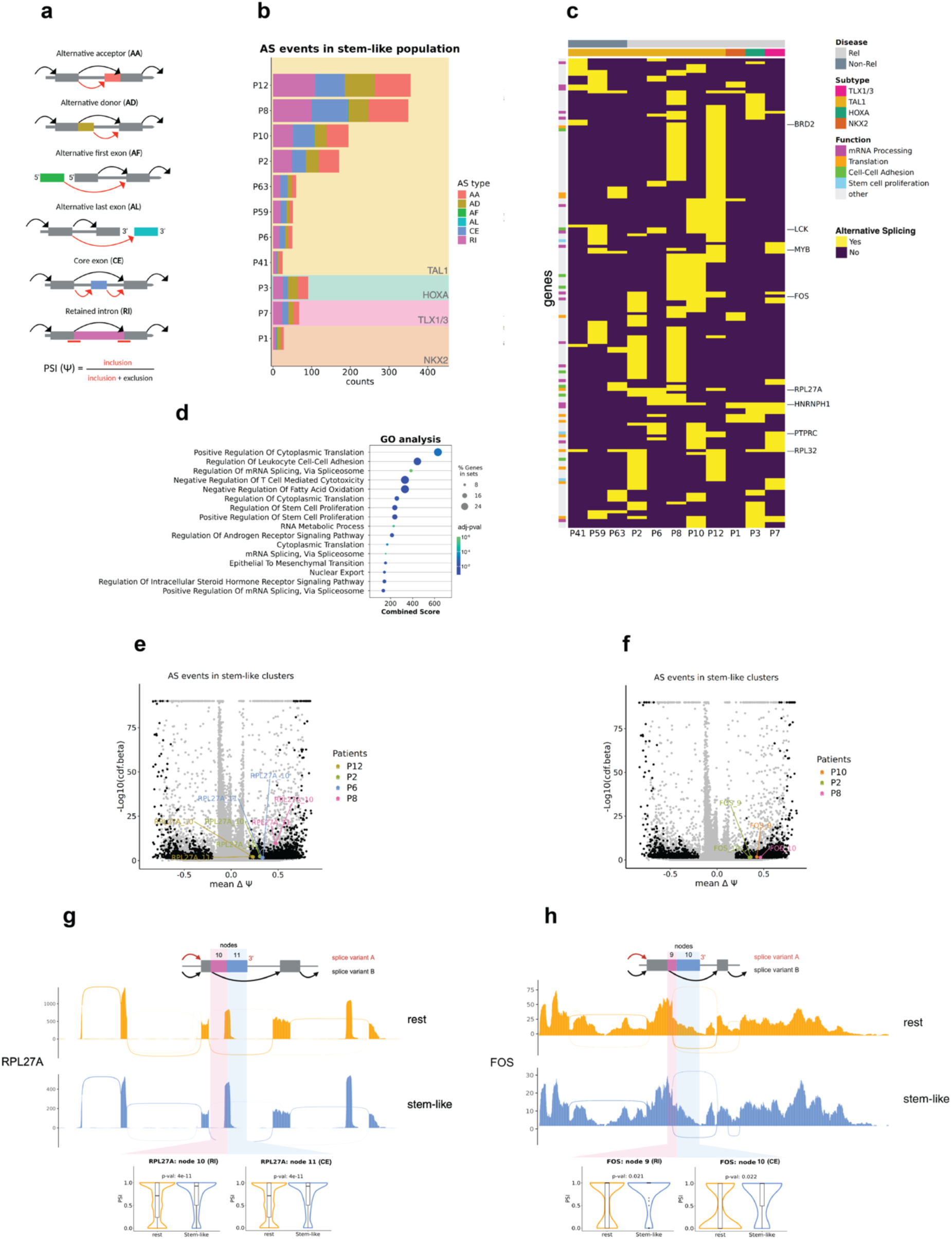
Alternative splicing variants are enriched in stem-like cell population. a) Illustration of all possible AS event types identified by the workflow (Suppl. Methods, Supplemental Figure 4a). Arrows indicate different possible splice junctions for each splicing node. Each color represents a different splicing event type. Psi (ψ) value refers to the number reads supporting the inclusion of the splicing node divided by the total number. Created with BioRender.com. b) Type and counts of detected AS events of individual T-ALL patients with >5% stem-like cell population. Background colors categorize patients according to their T- ALL subgroup. Partially created with BioRender.com. c) List of genes (rows) affected by alternative splicing in stem-like clusters of each patient (column). Yellow indicates an alternative splicing variant in the stem-like cell cluster of the patient, purple indicates no significant difference between stem-like cell cluster and other clusters of the patient. Patients are grouped based on their T-ALL subtype and by non-relapsing and relapsing disease. Left side of the panel highlights the biological pathways most affected by AS. Partially created with BioRender.com d) GO analysis represents significant enrichment of pathways (GO Biological Processes 2023) using all genes from c) as input. e), f): volcano plot illustrates inclusion rates (x-axis) of splicing nodes in stem-like cell cluster. Y axis represents significance values (-Log10(cdf.beta), see Suppl. Methods). All significant events are shown in black. e) volcano plot shows splicing events of four patients: P2, P6, P8, P12. Colors highlight two enriched splicing nodes (RPL27A node 10 and RPL27A node 11) in stem-like cell cluster shared among all four patients. f) volcano plot shows splicing events of three patients: P2, P8, P10. Colors highlight two enriched splicing nodes (FOS node 9 and FOS node 10) in stem-like cell cluster shared among all three patients. g), h): Upper panel: schematic representation of two splicing variants observed in illustrated splicing nodes. Pink node indicates a retained intron, blue node indicates a core exon (see illustration in a). Red arrow illustrates the splicing variant enriched in stem-like cells. Middle panel) Sashimi plots of stem-like cells (yellow) and other leukemic cells (blue) illustrate whole gene affected by alternative splicing. Read coverage is computed on P2, P6, P8, P12 (g) or P2, P8, P10 (h). Lower panel: Violinplot represents Psi (ψ) value of affected splicing nodes in stem-like cell cluster (yellow) vs other leukemic cells (blue). g) Affected splicing nodes of RPL27A gene. Partially created with BioRender.com. h) Affected splicing nodes of FOS gene.

To understand the biological processes in which AS occurs in these cells, we performed Gene Ontology (GO) based over-representation analyses, which revealed enrichment of AS genes in pathways presumed to contribute to the stemness phenotype (Figure 4d) - in close agreement with and thus corroborating our aforementioned gene expression analyses. These include cell adhesion (padj <0.01), stem-cell proliferation (padj < 0.05), and RNA metabolism (padj < 0.00001). In addition, as previously described in precursor B-ALL^31^, we find that splicing factors themselves are also affected by AS (padj < 0.000001).

We next elaborated on whether some of these genes are recurrently undergoing AS in T-ALL, (Figure 4c) implying they could potentially serve as a future treatment target. Indeed, we found 161 genes that are affected by AS in at least two patients. This includes known T-ALL oncogenes (e.g., LCK, MYB, FOS; detected in four patients) and tumor suppressors (e.g., PTPRC; detected in four patients), splicing factors (e.g., HNRNPH1; detected in six patients), ribosomal proteins (e.g., RPL32, RPL27A; detected in six and five patients) and epigenetic modifiers (e.g., BRD2; detected in three patients).

One of the recurrent events enriched in stem-like cells is a retained intron at node 10 and an additional exonic segment at node 11 of RPL27A which is shared among four patients (P2, P6, P8, P12) (Figure 4e,g). The retention of these two nodes results in a premature stop codon and in an open reading frame for a non-coding protein (ENST00000530585) which is a likely target of nonsense mediated decay (NMD)^34,35^. The finding of a likely inactivating alternative splicing event is compatible with the aforementioned downregulation of RNA translation and metabolism in the dormant population.

Next, we investigated the different splice isoforms of FOS, as FOS is one of the significant regulons in the stem-like cell population (Figure 4f,h). P2, P8 and P10 show a higher inclusion of node 10 and 11 in their stem-like cells which, interestingly, leads to a shortened but functional protein isoform (ENST00000554617^34^). While the consequences of this shortened version on the activity of FOS is currently unknown, the critical function of FOS in stem cell biology^36^ suggests a putative mechanism of mediating stem-like cell functions.

Together with the known impact of AS on the persistence of resistant subpopulations in hematologic malignancies^37^, these data suggest that specific splicing isoforms may contribute to stemness and treatment resistance in T-ALL, although the direct validation of potential mechanisms governing such function remain beyond the scope of this study.

### Analysis of T-ALL clonal evolution reveals marked enrichment of the stem-like cell population during relapse

Considering the dormancy phenotype and the downregulation of metabolic processes of the stem-like cells, we hypothesized that this population may be particularly treatment resistant. We therefore analyzed the clonal evolution from initial disease to relapse in those patients in whom the proportion of stem-like cells could be reliably quantified. We thus selected 8 out of the 13 patients in whom stem-like cells contribute to at least 5% of the total population (Figure 5). Stem-like cell clusters of each patient were identified based on the highest stemness score compared to other clusters of the same patient. At the time of initial diagnosis stem-like cells accounted for a very small proportion of all cells (0.46% - 2.16% in 6 patients) or remained undetectable in 2 patients. In contrast, by the time of relapse, the cells with a stem-like cell phenotype accounted for a substantial proportion of the total cells ranging from 10.68% to 43.52% (Figure 5d, paired t-test: p = 6.6e^−05^). Remarkably, four patients (P6, P1, P3, P7) display at least one more cluster exhibiting a high stemness score (cluster with more than 50% cells > 0.25) which is strongly enriched in the relapse sample. These data indicate that the stem-like cell population commonly expands during the transition from initial disease to relapse, likely reflecting resistance to first-line therapy. By contrast, the initially predominant subclones had largely disappeared in the relapse samples, which is consistent with the treatment response to first-line therapy largely resulting in low levels of residual disease or even transitory complete remission.

**Figure 5:**
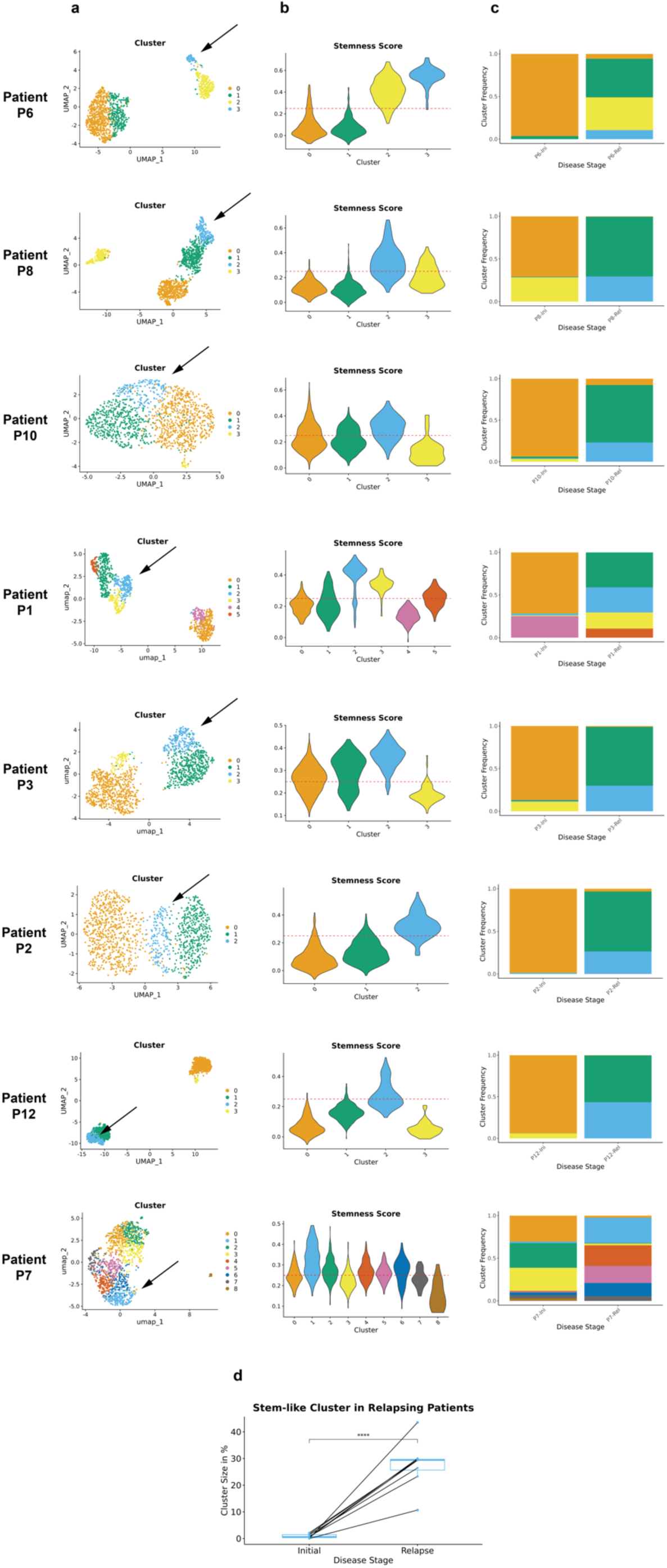
Stem-like cells expand during the transition from initial diagnosis to relapses. Data from 8 patients with >5% stem-like cells are shown. a) UMAPs based on RNA expression performed for individual patients. Black arrow points towards the cluster with the highest stemness score. b) Stemness score of clusters in individual patients displayed by violinplots. Red dashed line reflects the threshold used for the definition of stem-like cells (0.25) (see Figure 3c-d). c) Stacked barplots display the frequency of clusters at the time of initial diagnosis and relapse in individual patients. d) Boxplot displays significant enrichment in the frequency of stem-like cells at the relapse compared to initial diagnosis in all patients with more than 5% stem-like cells in total (paired t-test: p = 6.6e^−05^).

### The stem-like cells tolerate conventional treatment while highly proliferative cells remain sensitive in T-ALL relapse

We next sought to validate the hypothesis that the dormant cell population is less sensitive to treatment compared to the highly proliferative leukemia cells. We thus tested potential differences by treating relapse cells obtained from two patients (P1 and P6) *in vitro* for three days with Cytarabine (a key component of current treatment protocols). These cells were subsequently analyzed by high-throughput single-cell transcriptomics (10x Genomics scRNA-seq), resulting in 52,977 additionally sequenced single cells (Suppl. Methods). In the relapse sample of P1, the proportion of the cluster with the highest stemness score (Cluster 2, see Figure 5b) is significantly enriched (permutation test: FDR <0.05, log2FD >0.25) following treatment while the predominant population of cells grouping with cluster 1 (see Figure 5b) with a stemness score < 0.25 are largely sensitive to Cytarabine treatment (Figure 6a,c) (permutation test: FDR <0.05, log2FD <0.25). In the relapse sample of P6, two clusters (Cluster 2 and Cluster 3, see Figure 5b) mostly consist of cells with a stemness score > 0.25. Both of these clusters are significantly enriched upon Cytarabine treatment (permutation test: FDR <0.05, log2FD >0.25) while the remaining cells with a low stemness score (cells of Cluster 0 and Cluster 1, see b) are sensitive to treatment (Figure 6b,d) (permutation test: FDR <0.05, log2FD <0.25).

**Figure 6:**
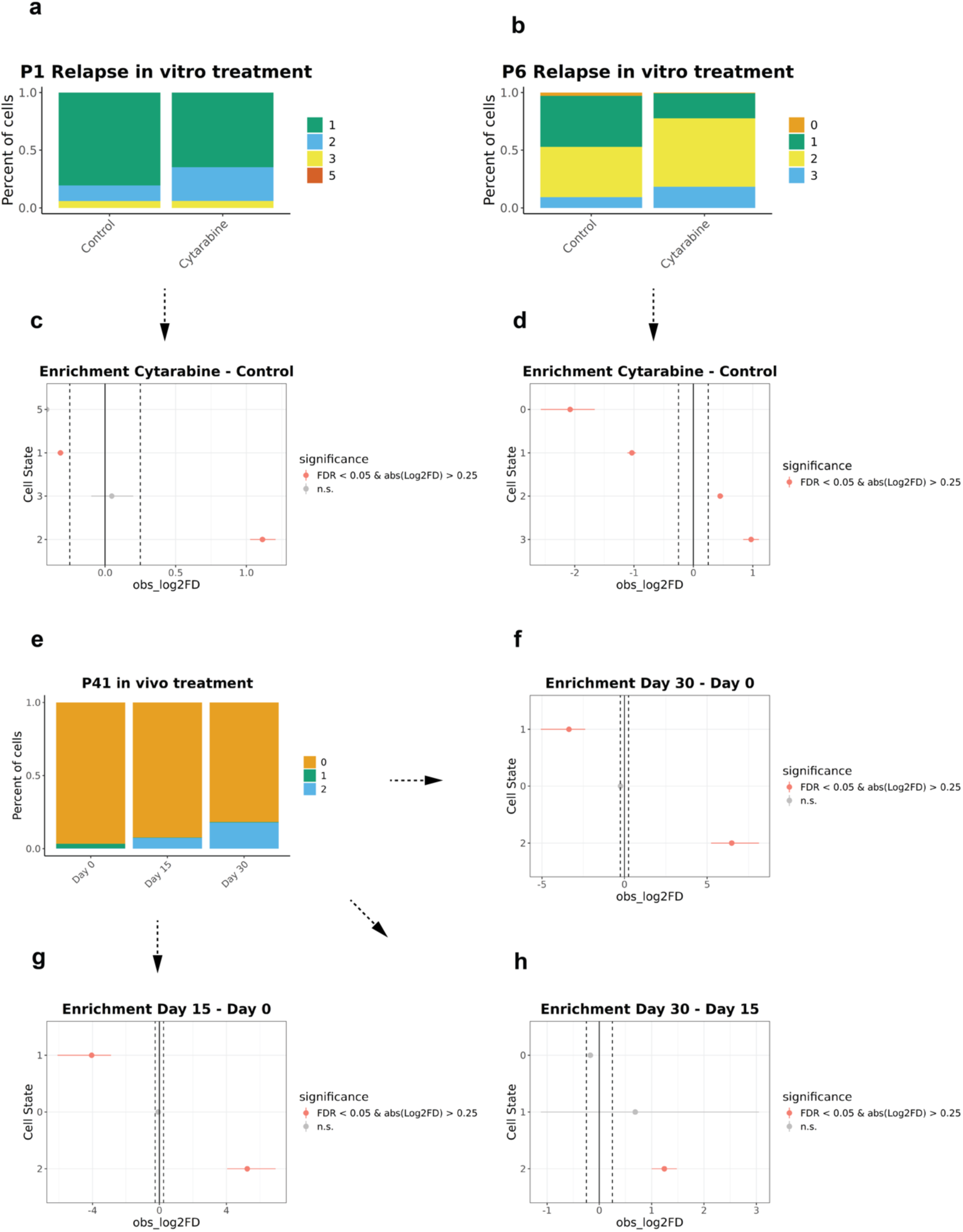
*In vitro* and *in vivo* treatment sensitivity of single cells. a), b): Stacked barplots display the frequency of clusters with DMSO control treatment (left) and 1μM Cytarabine treatment *in vitro* (right). c,) d): Analyses of differences in proportion of clusters between DMSO control and cytarabine *in vitro* treatment using permutation tests. N = 1000 permutations per analysis, significance threshold: FDR <0.05, log2FD > 0.25. a), d): relapse cells of P1. b), d): relapse cells of P6. e) Stacked barplots display the frequency of clusters in P41 on day 0 (left) and after 15 (middle) and 30 (right) days of treatment with a combination of Doxorubicin, Dexamethasone and Vincristine (right). f,) g), h): Analyses of differences in proportion of clusters between 0, 15 and 30 days of *in vivo* treatment using permutation tests. N = 1000 permutations per analysis, significance threshold: FDR <0.05, log2FD > 0.25.

To further validate these results, we analyzed T-ALL patient cells in a murine PDX *in vivo*. PDX cells of patient P41 (Supplemental Figure 5) were re-engrafted and, *in vivo*, treated with a combination of Vincristine, Doxorubicin and Dexamethasone (Suppl. Methods). After 15 and 30 days of treatment, samples were analyzed by scRNA-seq. The proportion of the stem-like cell cluster vastly increases after 15 days of treatment and enriches even further after 30 days (Figure 6e-h) (permutation test: FDR <0.05, log2FD >0.25), validating the treatment resistance of cells with a high stemness score *in vivo*. By contrast, the proportion of cells with a low stemness score is reduced following treatment confirming that this type of cells is sensitive to chemotherapy.

## Discussion

Understanding the mechanisms driving relapse and treatment resistance in T-ALL remains an unmet clinical need. While stem-like cells have been discussed as a potential type of cells with particular chemoresistance, the cellular networks underlying such resistance can only be incompletely explored by bulk sequencing. This limitation is aggravated, as stem-like T-ALL cells presently cannot be readily identified and isolated by standard techniques.

Harnessing VASA-seq^8^, we provide comprehensive insights into the transcriptomic landscape of T-ALL from initial diagnosis to relapse, by unveiling a previously elusive subpopulation of T-ALL cells characterized by a quiescent, stem-like cell phenotype. We identified distinct transcriptional signatures, gene regulatory networks and splicing isoforms which we hypothesize to govern the biology and the treatment resistance of these stem-like cells.

Consistent with previous findings in precursor B-ALL, a large fraction of stem-like cells persisted in G1 phase and upregulated stem-cell related pathways, such as cell adhesion facilitating the interaction with the stromal environment and resulting in the protection against external factors in the stem-cell niche^13,38^. Moreover, the analysis of recurrent AS revealed an enrichment in such cells in mRNA isoforms potentially impacting the metabolic activity, including those encoding ribosomal proteins (such as RPL27A), and those that are expected to modulate stem-cell induction, including FOS^36^. Indeed, AS might provide a post-transcriptional source for the regulation of stemness networks in T-ALL.

Mapping the expression profiles of the T-ALL samples to a thymic reference^11^ assigned the majority of stem-like cells as DN progenitors. Our data thus indicate that T-ALL stem-like cells are particularly immature, which is consistent with the exquisite treatment resistance of this type of cells.

Remarkably, our data imply that the clonal expansion of stem-like cells is much more common in TAL1-driven T-ALLs than in the other subgroups. Considering that most type-2 T-ALL relapses are TAL1-driven^3,5^, these findings indicate that these early leukemic ancestors may include stem-like cells as defined in our study. This also aligns with the hypothesis that stem-like cells can derive from pre-leukemic clones in early phases of the disease^12,39,40^. Nonetheless, we found stem-like cells also in HOXA-, NKX2-, and TLX1/3- driven T-ALLs that mostly relapse as type-1, albeit at lower frequencies. Importantly, the prediction of treatment resistance was functionally validated by *in vitro* and *in vivo* drug-testing thus identifying a defined cell population that accounts for only a minority of the leukemic cells at initial diagnosis but has expanded substantially by the time of relapse.

Given the need for developing new treatment strategies, the stem-like cells defined in our study should be considered as future therapy targets to prevent relapse and overcome resistance. The stemness score developed in our study may, thereby, identify this resistant cell population, serving as a clinical biomarker.

## Acknowledgments

We thank all the patients who participated in the study and their families. We thank EMBL FCCF Services, EMBL IT Services and EMBL GeneCore for technical support. We further thank D. Campana, St. Jude Childreńs Research Hospital, Memphis, TN, for kindly providing MSCs. This work was supported by the Athenaeum Dietrich Götze Stiftung für Kultur und Wissenschaft.

## Authorship

### Contribution

JC designed and performed the scRNA-seq experiments, run bioinformatic analyses and wrote the manuscript. KKR established PDX models. KKR performed drug treatment assays. PZ performed splicing variant analysis and run SCENIC pipeline. TR supervised computational analysis. AM helped with the set-up of the SCENIC pipeline^14,15,41^ and run initial SCENIC analyses. JZ supervised the set-up of the SCENIC pipeline^14,15,41^. MS, GE, CE provided patients samples and data for the analyses. JPB and BBR supervised the establishment of the PDX models and drug treatment assays. AEK and JOK designed the research, supervised the project, and wrote the manuscript. All authors reviewed and contributed to the final manuscript.

