## Supplementary Materials for "Role of Stem-Like Cells in Chemotherapy Resistance and Relapse in pediatric T Cell Acute Lymphoblastic Leukemia"

### Supplemental Methods

#### Patient-derived xenografts (PDXs)

We maintained T-ALL patient cells by intrafemoral injection of $1 \times 10^5$ to $5 \times 10^6$ viable primary ALL cells in NSG (NOD.Cg-PrkdscidIl2rgtm1Wjl/SzJ) mice. Transplanted mice were both male and female, aged 5–8 weeks. Animals were housed in individually ventilated cages with access to food and water ad libitum. Leukemia progression was monitored in the peripheral blood by flow cytometry using anti-mCD45, anti-hCD45, anti-hCD19, or anti-hCD7 antibodies. Cells were harvested after engraftment had reached 75% in the peripheral blood or the mice health score had reached either three at single item or the total score had reached five. T-ALL cells were collected from spleen and cryopreserved as described[1]. Leukemia progression was monitored in the peripheral blood by flow cytometry using anti-mouse-CD45 (anti-mCD45), anti-human-CD45 (anti-hCD45), anti-hCD19, or anti-hCD7 antibodies. Xenograft identity was verified by DNA fingerprinting using the commercial AmpFlSTR® NGM Select kit.

#### Primary cell culture

Human hTERT immortalized primary bone marrow mesenchymal stroma cells[2] (MSC)were cultured in RPMI 1640 medium supplemented with 10% heat-inactivated fetal bovine serum, L-glutamine (2 mM), penicillin/streptomycin (100 IU/ml) and hydrocortisone (1 µM).

#### scRNA-seq: library preparation

Cryopreserved cells were thawed at 37°C and resuspended in 10 ml RPMI medium + 20 % FBS. Cells were centrifuged for 5 min at 300g and resuspended in ice-cold PBS + 2% FBS + 5mM EDTA. Cells were stained on ice and in the dark for 30 min with anti-murine-CD45-PE (mCD45)(clone 30-F11; BioLegend; 1:20). 1:100 4,6-diamidino-2-phenylindole (DAPI) was added immediately before FACS sorting. mCD45-DAPI- cells were sorted using a BD FACSAria fusion cell sorter.

For VASA-seq, single cells were sorted into cooled 384-well plates containing primers with well-specific barcodes. After sorting, plates were immediately spun and placed on dry ice. Plates were shipped on dry ice to Single Cell Discoveries (SCD), where libraries were prepared by the company according to the VASA-seq protocol[3] and sequenced on the Nova Seq X Plus (10B - 100 cycle, paired-end, 150.000 reads/cell). Mapping, *in silico* depletion of rRNA and generation of count tables were automated using the STARSolo aligner by SCD. For 10x Genomics bulk of cells was sorted in ice-cold PBS + 3% BSA. Samples were processed immediately according to standard 10x Genomics Chromium 3′ (v.3.1 Chemistry) protocol. Libraries were sequenced on the NovaSeq 6000 S2 v1.5 (100 cycles) flowcell S2 (Surface: 3.3-4.1 BIO reads). Transcripts were quantified into count matrices using cell ranger mkfastq and count workflows (10x Genomics, v.3.1.0, default parameters). Sequenced transcripts were aligned to a mixed human-mouse genome (GRCh38-GRCm38). The R package Seurat was used for downstream analysis[4].

#### scRNA-seq: Data preprocessing and quality control

Human cells were separated from mouse contaminations by filtering for cells with more than 80% of transcripts aligning to GRCh38. Remaining reads aligned to the mouse genome were removed from the analysis.  Filtering of low-quality cells based on gene count (VASA-seq:

>2500 and <9000, 10x Genomics: >300 and <7500), read count (VASA-seq: <100000, 10x Genomics: <40000) and mitochondrial fraction (VASA-seq: <5%, 10x Genomics: <10%) was performed.

**scRNA-seq: Dimensionality reduction and clustering based on RNA expression**

Downstream processing involved log2normalization, scaling and linear dimensional reduction (PCA). We assigned cell cycle scores using the CellCycleScoring() function of Seurat and canonical histone expression scores using the AddModuleScore() function. UMAP reduction and differential expression analysis of identified clusters was performed un uncorrected samples following the standard Seurat workflow. For the integrated analysis of patients (see Supplemental Figure 2c), samples were batch corrected using the anchor-based CCA integration of Seurat. For gene expression analyses of clusters, the FindMarkers() function was applied[4]. Gene-Set Enrichment Analysis (GSEA) was performed using the prerank() module from the gseapy Python package with standard parameters[5]. The Normalized Enrichment Score (NES) facilitates the comparison of analysis results across different gene sets by normalizing to the average Enrichment Score (ES) from all dataset permutations. To infer cell types, a thymic single cell atlas was used as a reference[6] and mapped onto our dataset using the functions FindTransferAnchors()and MapQuery() of the Seurat package[4]. To infer clustering on the drug and control treated 10x Genomics data based on the respective untreated VASA-seq samples, the latter were used as a reference[6] and the 10x Genomics data were mapped onto it using the functions FindTransferAnchors() and MapQuery() of the Seurat package[4].

**scRNA-seq: Gene-regulatory networks and dimensional reduction based on regulon activity**

Gene-regulatory networks of VASA-seq data was inferred using an in-house constructed Snakemake pipeline[7] of the SCENIC package[8,9]. SCENIC calculates regulon activity in three steps: 1) generation of co-expression modules of transcription factors and genes (GENIE3). 2) removal of indirect targets using TF motif information (RcisTarget). 3) calculation of regulon activity in individual cells (AUCell). Scaling and UMAP reduction were performed on the uncorrected patient samples using the regulon activity. For differential expression, cells were merged into two larger clusters (Stem_like and Blasts) to focus on specific stem cell-like markers compared to other leukemic cells. The stemness signature of T-ALL cells was calculated with the AddModuleScore() function of Seurat using marker genes with a log2FC >0.5 and padj <0.05 as input for the *features* parameter[4]. Natural breaks in the stemness scores were identified by getJenksBreaks (k=4). Gene-Set Enrichment Analysis (GSEA) was performed using the prerank() module from the gseapy Python package with standard parameters[5]. The Normalized Enrichment Score (NES) facilitates the comparison of analysis results across different gene sets by normalizing to the average Enrichment Score (ES) from all dataset permutations. Gene sets were retrieved from the 'GO Biological Processes 2023' library.

**Alternative Splicing (AS) Analysis**

First, reads were demultiplexed and trimmed with TrimGalore (https://github.com/FelixKrueger/TrimGalore). Then, the barcode-specific FASTQs were ribo-depleted using both mouse and human ribosomal-DNA sequences as reference. Reads mapping uniquely to the mouse genome (GRCm38) were eventually discarded with BBMap (https://sourceforge.net/projects/bbmap/). The quantification and differential testing of AS events were performed by applying the specialized computational workflow published in Salmen et al[3]. Firstly, we expanded the transcriptome of each patient. In synthesis, FASTQ pseudo-bulks, corresponding to the previously identified clusters, were aligned to the reference human genome (GRCh38) via HISAT2 with standard configuration[27]**.** The resulting transcriptomes were assembled and merged with StringTie2[10]. The newly annotated isoforms underwent several quality-control steps to filter out possible false positives (see Methods in Salmen et al[3]). Consequently, high-confidence novel isoforms were added to the reference GTF file (release GRCh38.110).The successive step consisted in further expanding the transcriptome by adding novel microexons (exons < 30 nt) to the reference GTF annotation. Therefore, we employed the discovery module from MicroExonator, a Snakemake workflow specifically designed to identify and quantify microexons[11]. The filtering of spurious hits was performed following the authors' guidelines. To quantify AS events across cell clusters, we ran the MicroExonator's downstream module 'snakepool' with default parameters. Cells from the same cluster were randomly pooled together into 5 pseudo-bulks of equal size. The PSI (percentage-spliced-in) value returned for each splicing node in the pseudo-bulks was used to provide a probability of differential inclusion. To avoid false positives, the pseudo-bulk quantification and assessment of differential inclusion were repeated 50 times for each pairwise comparison. The probabilities of each splicing node were then fitted to a beta distribution and the CDF-beta value was returned. Splicing nodes with CDF-beta $< 0.05$, mean probability $> 0.9$, and $\Delta PSI > 0.2$ were considered differentially included, while events found significant in less than 25 repetitions were discarded. We designed the pairwise comparisons to expose the different AS patterns either between two distinct clusters or between one cluster and the rest of the sample.To quantify the splicing events at the single-cell level, we instructed MicroExontor to bypass the pseudo-bulk pooling and differential testing. Sashimi plots of specific AS events were created with the ggsashimi package[12].Gene Ontology (GO) over-representation analysis was performed using the enrichr() module from the gseapy Python package with standard parameters. Gene sets were retrieved from the 'GO Biological Processes 2023' library.

**Supplemental Tables Legend**

Supplemental Table 1: **Differential expression markers of individual Patient clusters**

Supplemental Table 2: **Differential expression markers of TAL1 stem-like cells vs other leukemic blasts**

Supplemental Table 3: **Differential regulon activity (SCENIC) of TAL1 stem-like cells vs other leukemic blasts**

Supplemental Table 4: **Differential regulon activity (SCENIC) between individual patients**

Supplemental Table 5: **Metadata and list of events of AS analysis**

**Supplemental Figures Legend**

Supplemental Figure 1: **Analyses of differences in cell cycle phases and predicted cell types using permutation tests**. N = 1000 permutations per analysis, significance threshold: FDR <0.05, log2FD > 0.25. a), b), c), e), f), g): analysis of patient P2. a) enrichment of predicted cell types in cluster 2 vs 0. b) enrichment of predicted cell types in cluster 1 vs 0. c) enrichment of predicted cell types in cluster 2 vs 1. e) enrichment of cell cycle phases in cluster 2 vs 0. f) enrichment of cell cycle phases in cluster 1 vs 0. g) enrichment of cell cycle phases in cluster 2 vs 1. d), h): analysis of TAL1 patients. d) enrichment of predicted cell types in stem-like cells vs other leukemic blasts. h) enrichment of cell cycle phases in stem-like cells vs other leukemic blasts.

Supplemental Figure 2: **Differential expression analysis, UMAP reductions and GSEA plots**. a) Heatmaps displays top50 enriched genes in in stem-like cells vs other leukemic blasts (log2FC >0.25, padj <0.05). Each column represents a single cell, each row a gene. b) UMAP reduction of TAL1-driven patients by RNA expression only. c) UMAP reduction of TAL1-driven patients by RNA expression after anchor-based CCA integration. d) UMAP reduction by SCENIC including patients of all T-ALL subgroups. e), f), g): GSEA plots of published datasets. e) signature of T cell quiescence genes (human orthologs were used)[13].f) signature of *de novo* prednisone resistance in T-ALL[14]. g) signature of T-ALL MRD[15].

Supplemental Figure 3: **Analyses of differences in the proportion of stem-like cells among T-ALL subgroups using permutation tests**. N = 1000 permutations per analysis, significance threshold: FDR <0.05, log2FD > 0.25.a) enrichment of stem-like cells in HOXA vs TLX1 cells. b) enrichment of stem-like cells in NKX2 vs HOXA cells. c) enrichment of stem-like cells in NKX2 vs TLX1 cells. d) enrichment of stem-like cells in TAL1 vs HOXA cells. e) enrichment of stem-like cells in TAL1 vs TLX1 cells. f) enrichment of stem-like cells in TAL1 vs NKX2 cells.

Supplemental Figure 4: **Computational workflow of AS analysis and single-cell quantification**. a) Representation of the four contiguous modules composing the workflow for the detection of AS in the stem-like cell population. First, raw FASTQ are pre-processed and mouse-reads discarded, then the transcriptome annotation is expanded with novel isoforms and microexons inferred from the data. The last module quantifies inclusion rates ($\psi$) at the single-cell level and performs statistical tests to identify AS events with high confidence. Module I, II, III were adapted from Salmen et al[3]. Created with BioRender.com. b) Representation of 'splicing node', the unit of measure of the AS workflow. For each gene, UTRs, exons (dark grey), and introns (light grey) from different isoforms are collapsed into unique and non-overlapping gene intervals, denominated splicing nodes. In addition to intron/exon junctions, splicing node boundaries are also inferred from the read coverage (e.g., node 3). Created with BioRender.com. c), d): $\psi$ UMAPs of individual patients. c) UMAPs of cells from P12, P8, P2, and P6 on which the $\psi$ value of RPL27A node 11 (left) and 10 (right) is projected. d) UMAPS of cells from P8, P2, P10 on which the $\psi$ value of FOS node 9 (left) and 10 (right) is projected. (c,d) Yellow is complete node inclusion ($\psi$=1), while blue is complete node exclusion ($\psi$=0). Gray cells lack information about the node.

Supplemental Figure 5: **Clonal composition of untreated P41**. a) UMAPs based on RNA expression performed for P41. b) Stemness score of clusters in P41 displayed by violinplots. Score is calculated as described in Figure 3c-d. Red dashed line reflects the threshold used for the definition of stem-like cells (0.25). c) Stacked barplots display the frequency of clusters at initial diagnosis.

### Supplemental Figures

Supplemental Figure 1

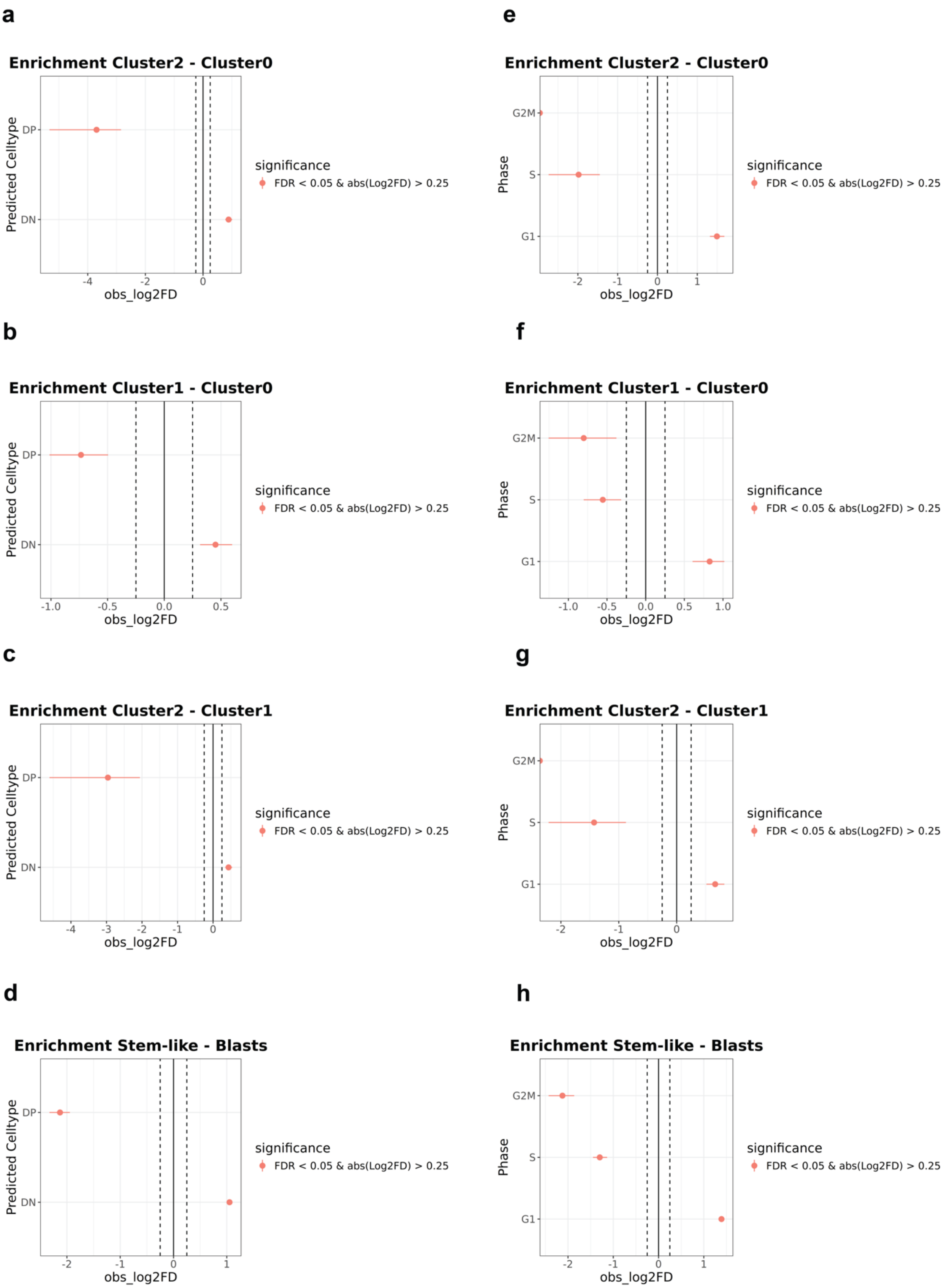

Supplemental Figure 2

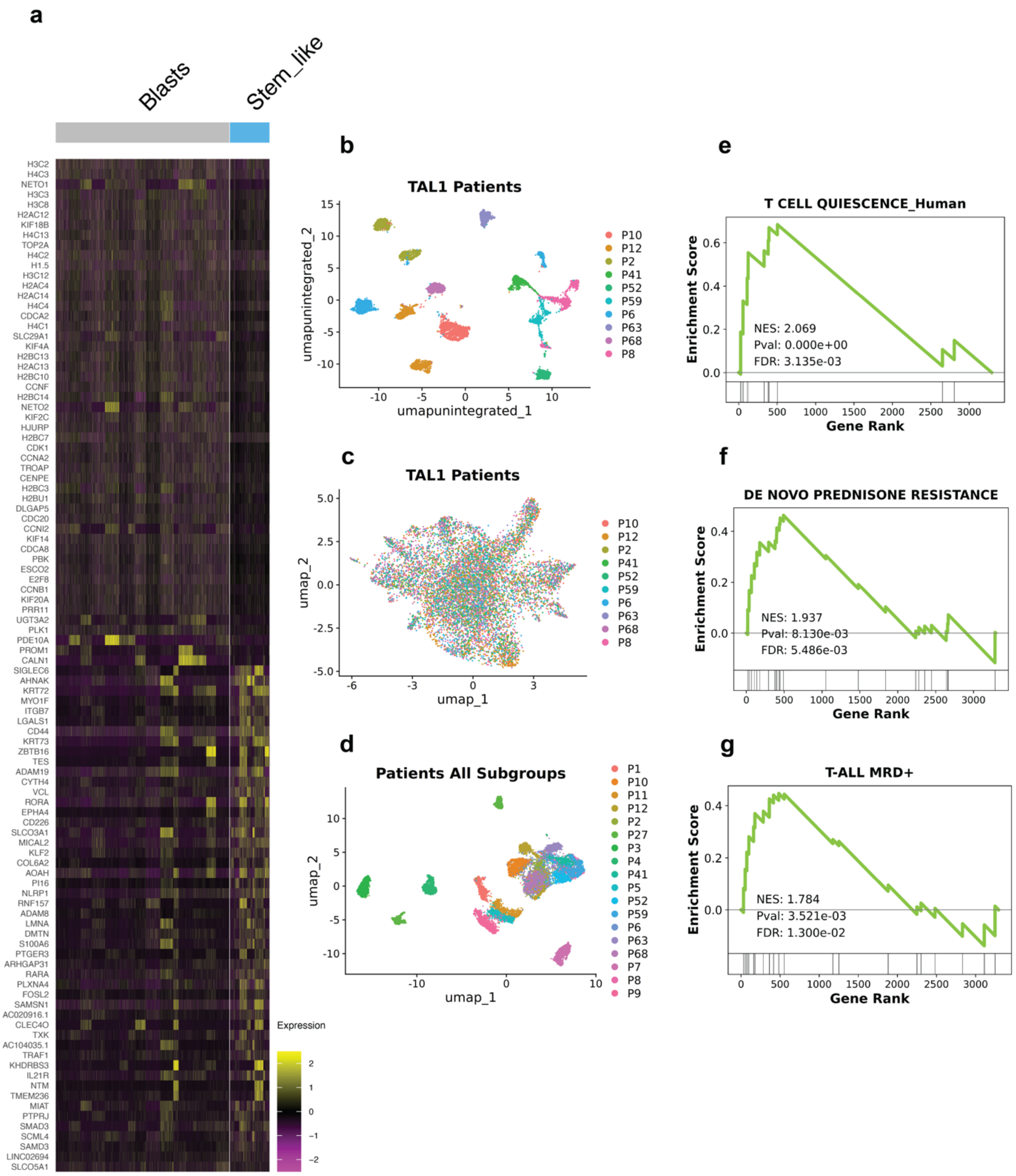

Supplemental Figure 3

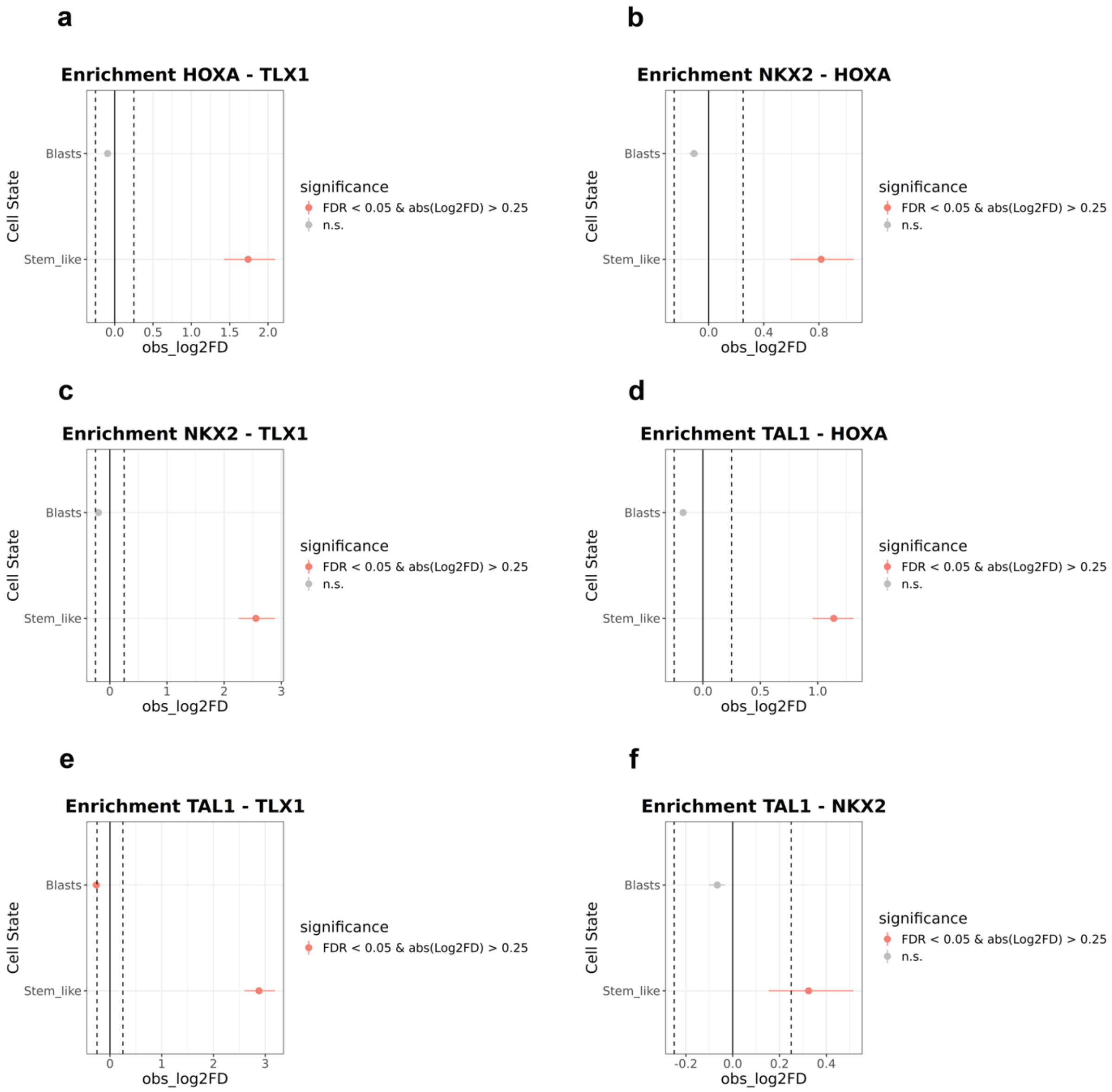

Supplemental Figure 4

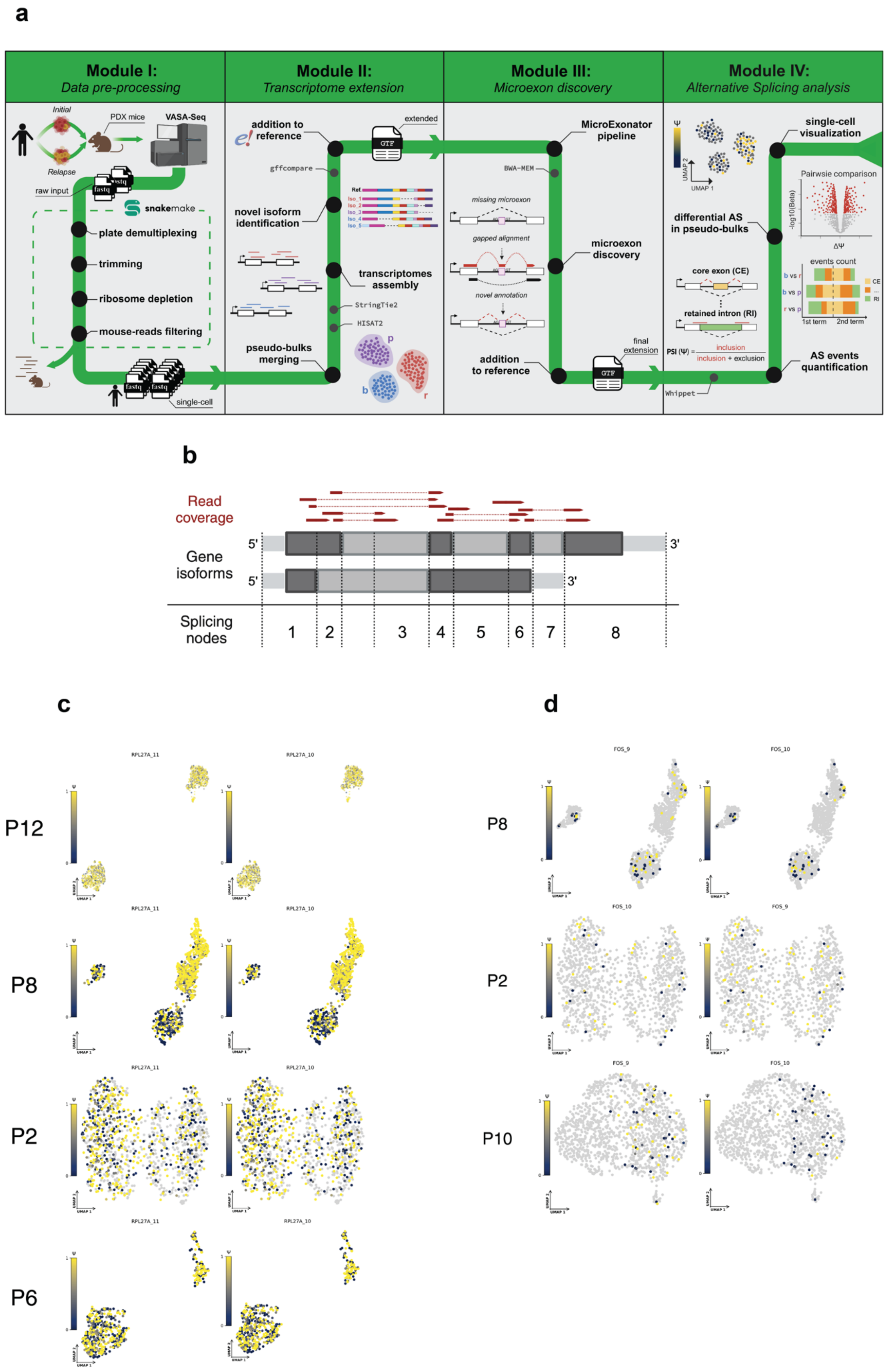

Supplemental Figure 5

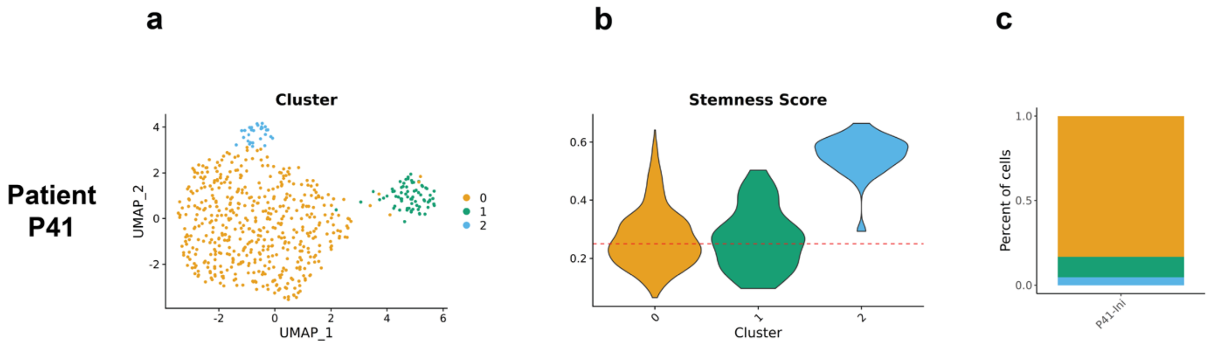
